## Supplementary material for "Graph-based pangenomics maximizes genotyping density and reveals structural impacts on fungal resistance": All supplemental files are located in this compressed directory and are described by manuscript name and file name in legend: FOM2 PCR.pptx

### Slide 1
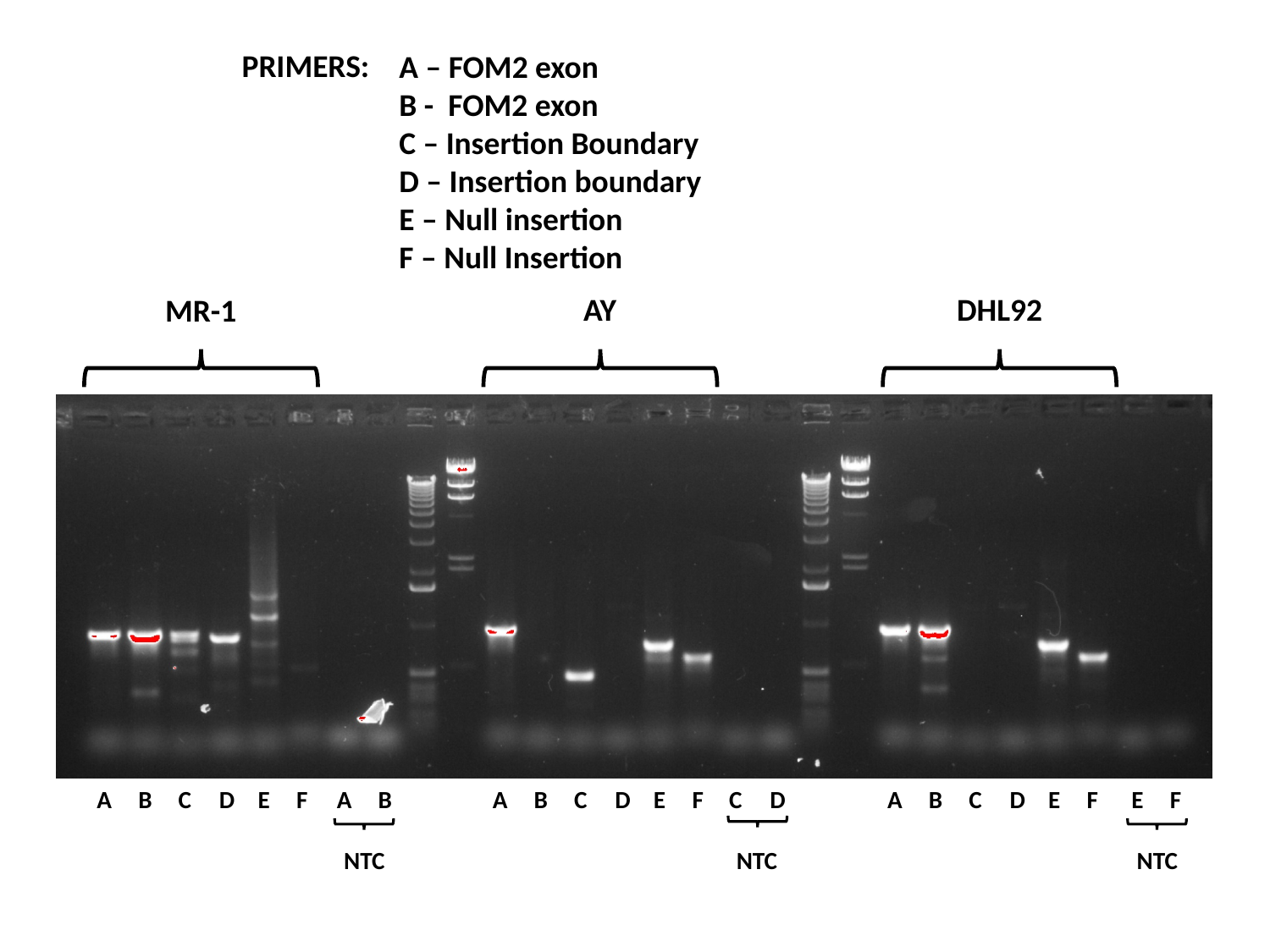

PRIMERS:
A – FOM2 exon
B - FOM2 exon
C – Insertion Boundary
D – Insertion boundary
E – Null insertion
F – Null Insertion
AY
DHL92
MR-1
F
C
E
B
D
F
A
F
F
E
E
E
C
C
C
D
D
D
B
B
B
A
A
A
NTC
NTC
NTC
