## Supplementary material for "Graph-based pangenomics maximizes genotyping density and reveals structural impacts on fungal resistance": All supplemental files are located in this compressed directory and are described by manuscript name and file name in legend: fom2_haplotypes_pcr.pdf

PRIMERS:
A – FOM2 exon
B - FOM2 exon
C – Insertion Boundary
D – Insertion boundary
E – Null insertion
F – Null Insertion

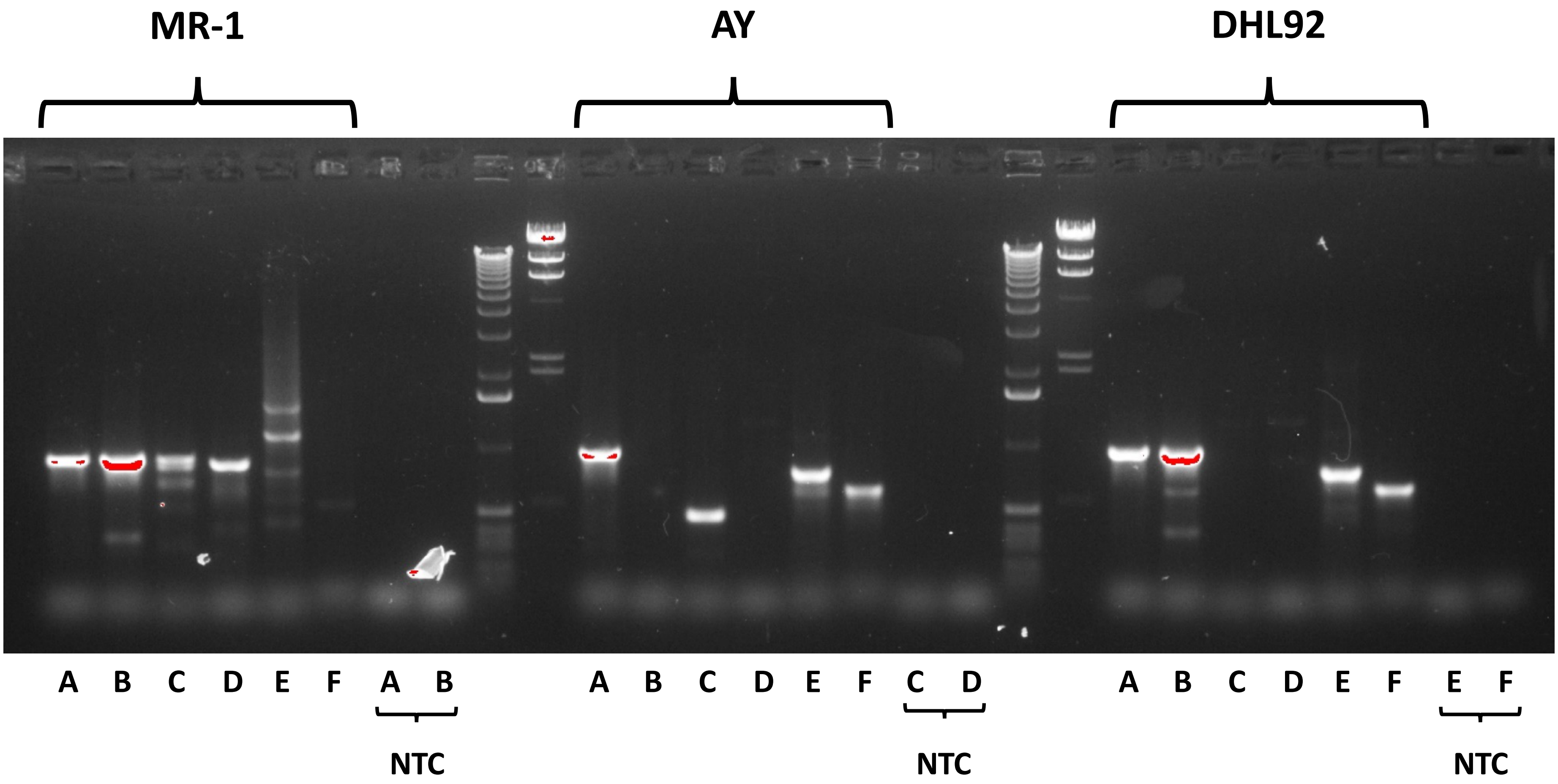
