## Supplementary material for "Graph-based pangenomics maximizes genotyping density and reveals structural impacts on fungal resistance": All supplemental files are located in this compressed directory and are described by manuscript name and file name in legend: powderyMildew.mixedModel.chromosomewise.pdf

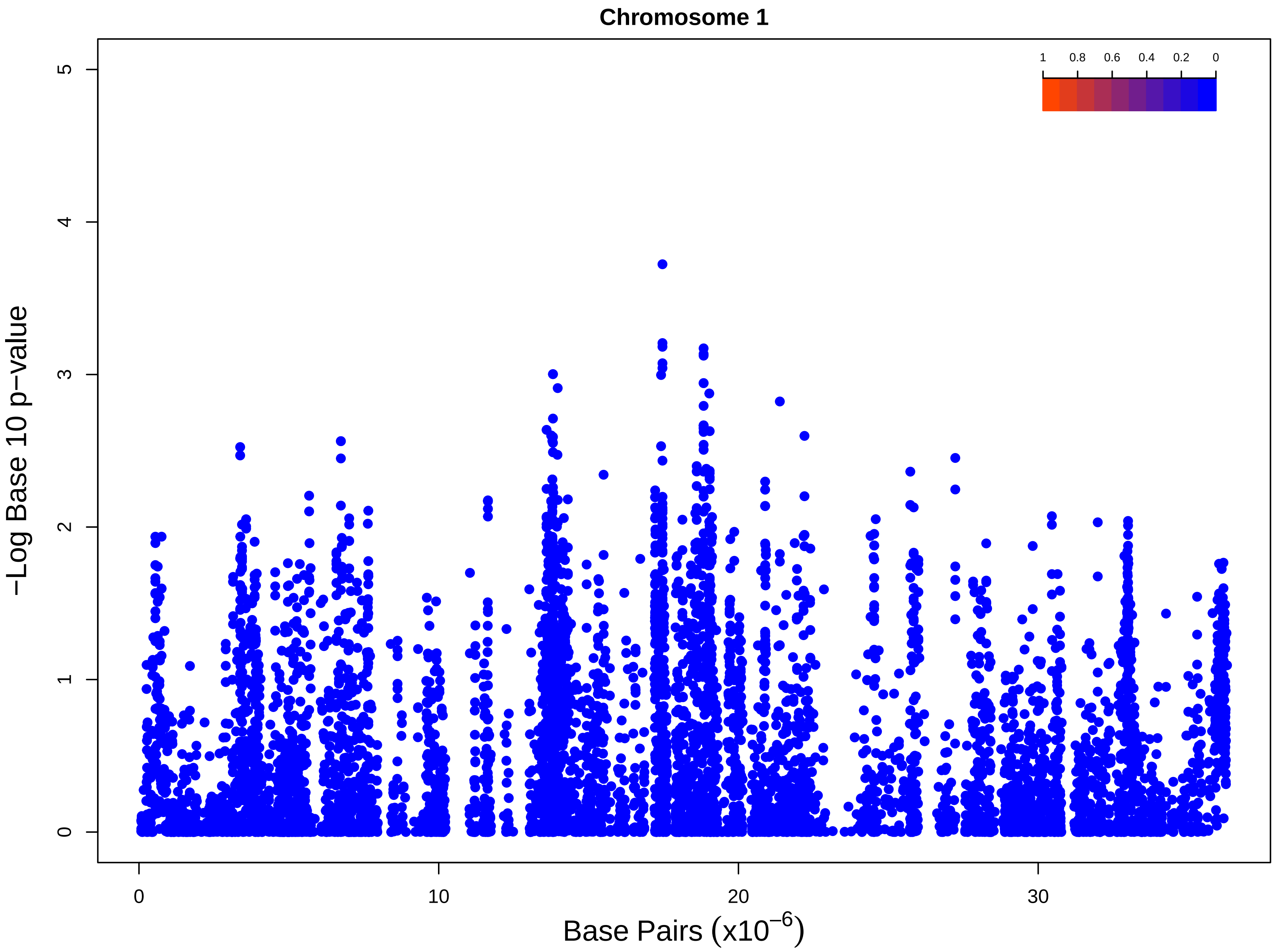
Chromosome 1
1 0.8 0.6 0.4 0.2 0
−Log Base 10 p−value
0 1 2 3 4 5
0 10 20 30
Base Pairs (x10$^{-6}$)

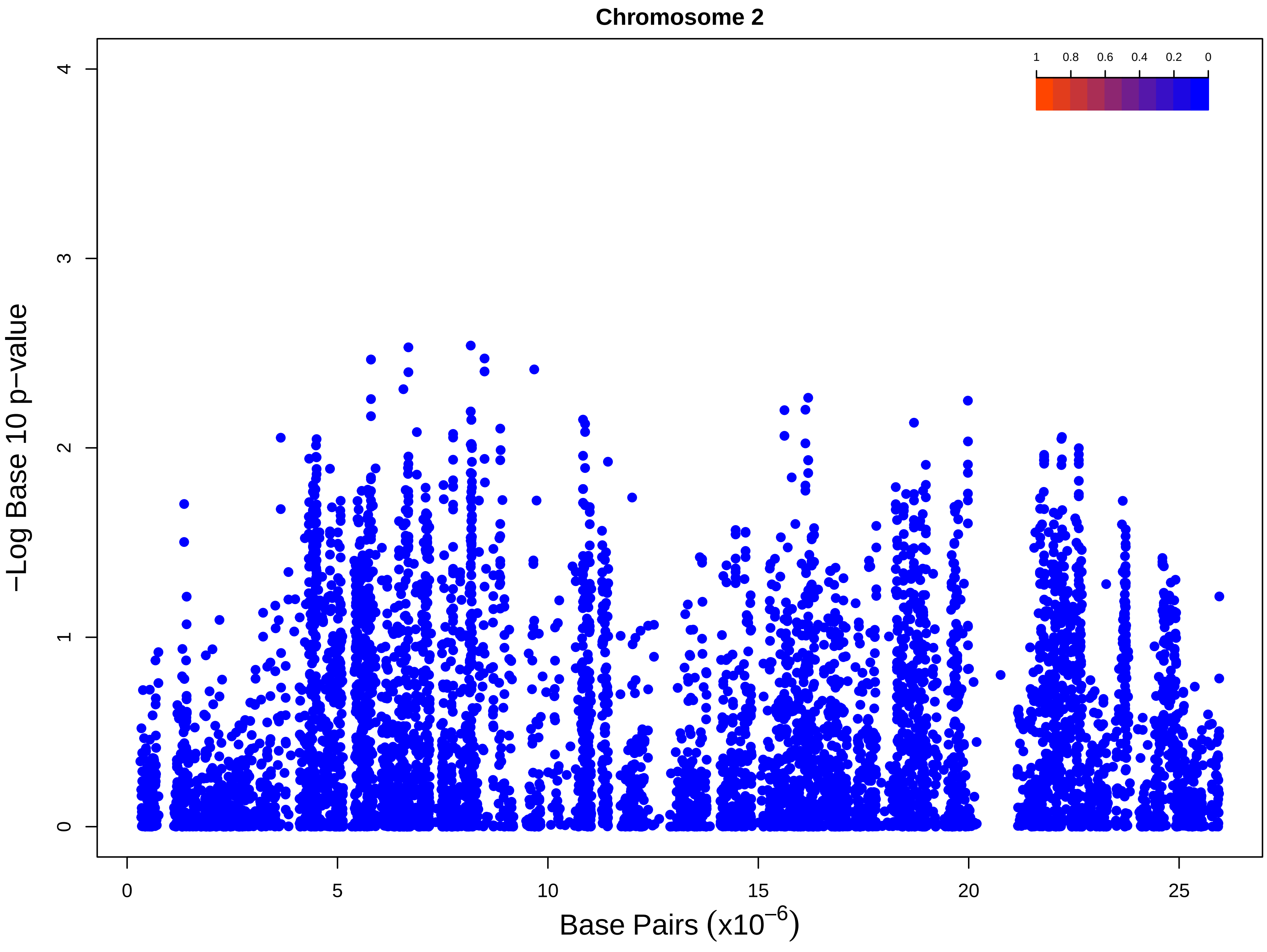
Chromosome 2
1
0.8
0.6
0.4
0.2
0
−Log Base 10 p−value
0
1
2
3
4
0
5
10
15
20
25
Base Pairs (x10$^{-6}$)

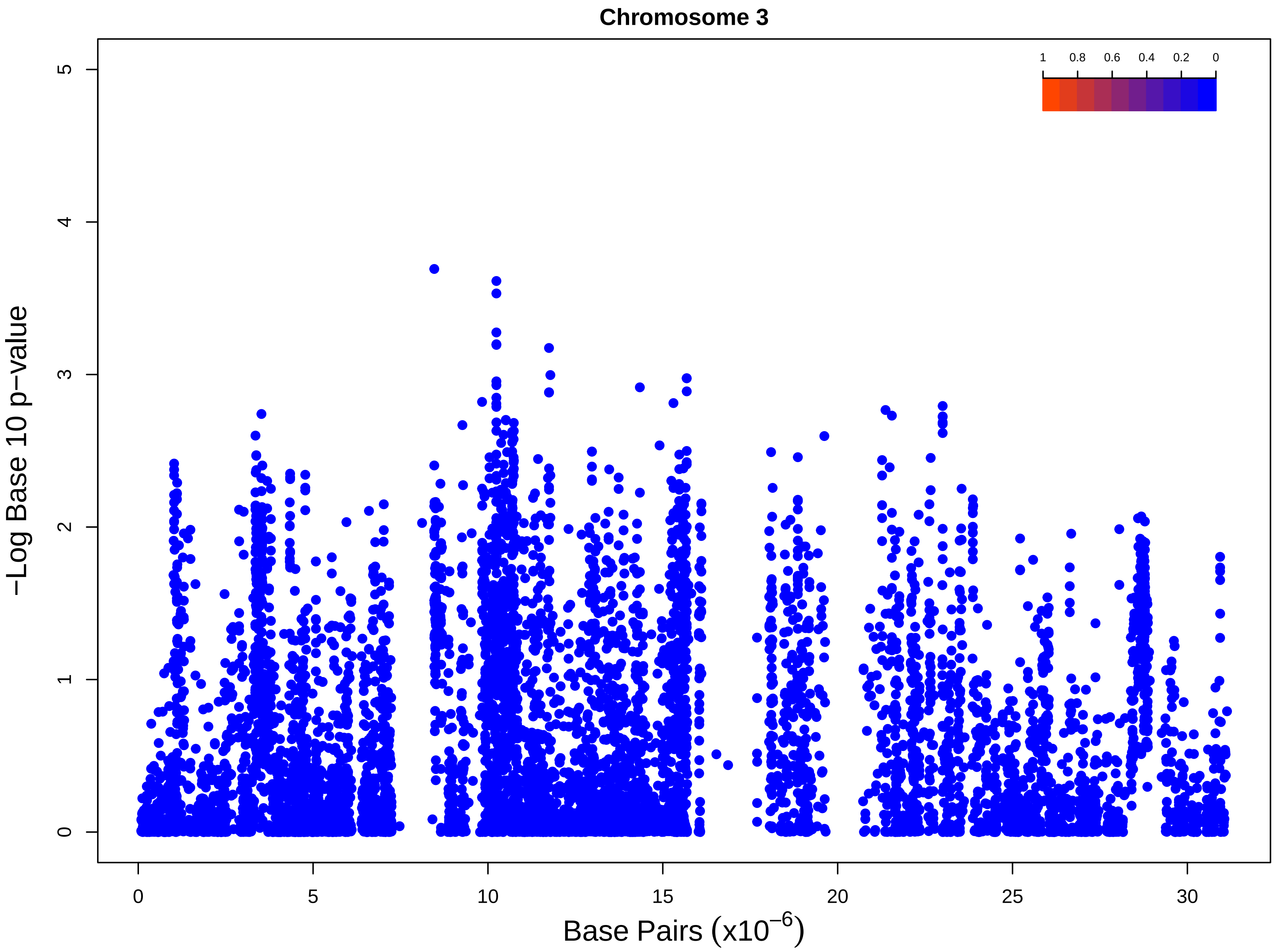
Chromosome 3
1
0.8
0.6
0.4
0.2
0
−Log Base 10 p−value
0
1
2
3
4
5
0
5
10
15
20
25
30
Base Pairs (x10^−6)

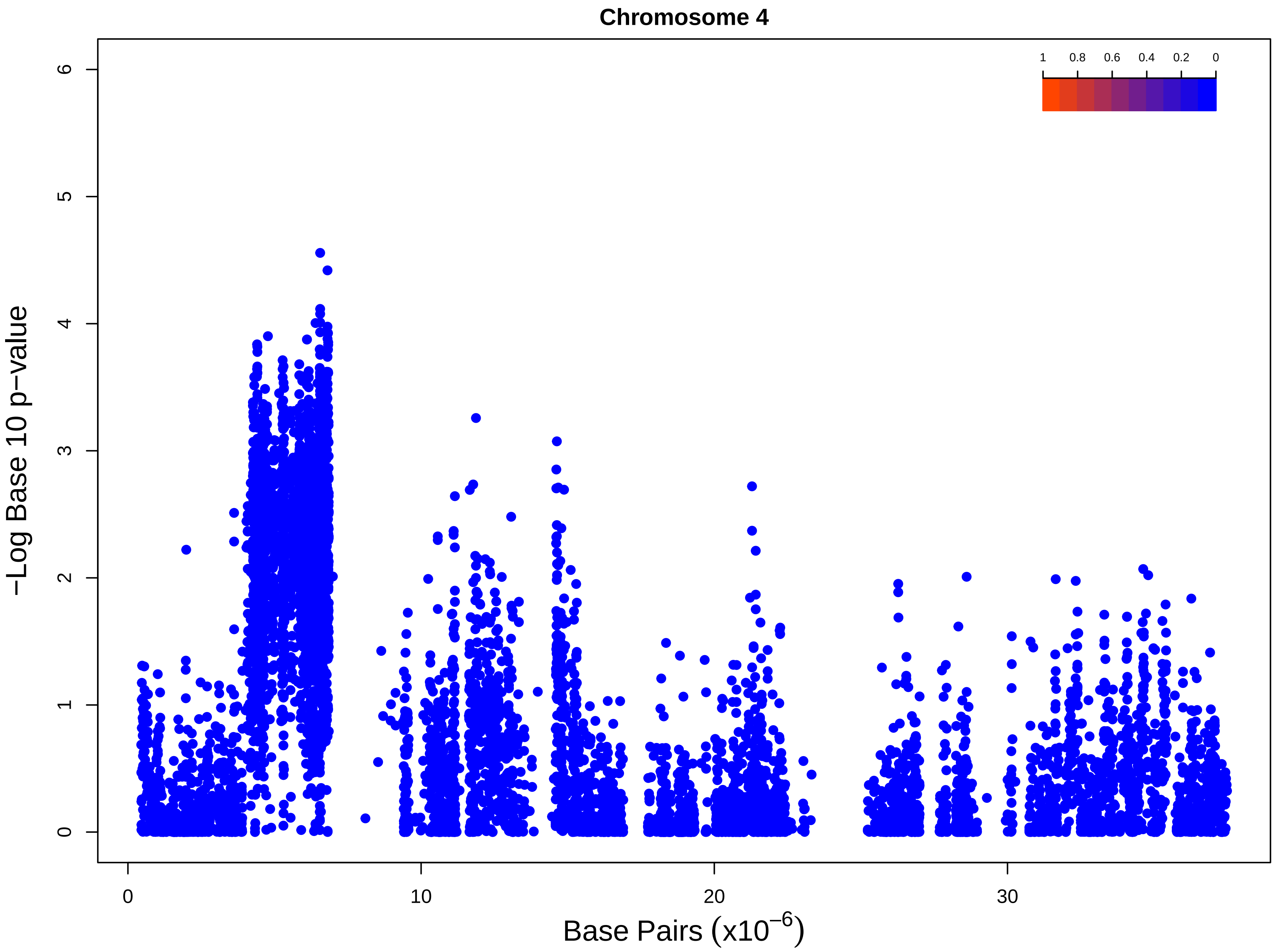

Chromosome 4
1
0.8
0.6
0.4
0.2
0
−Log Base 10 p−value
0
1
2
3
4
5
6
0
10
20
30
Base Pairs (x10^−6)

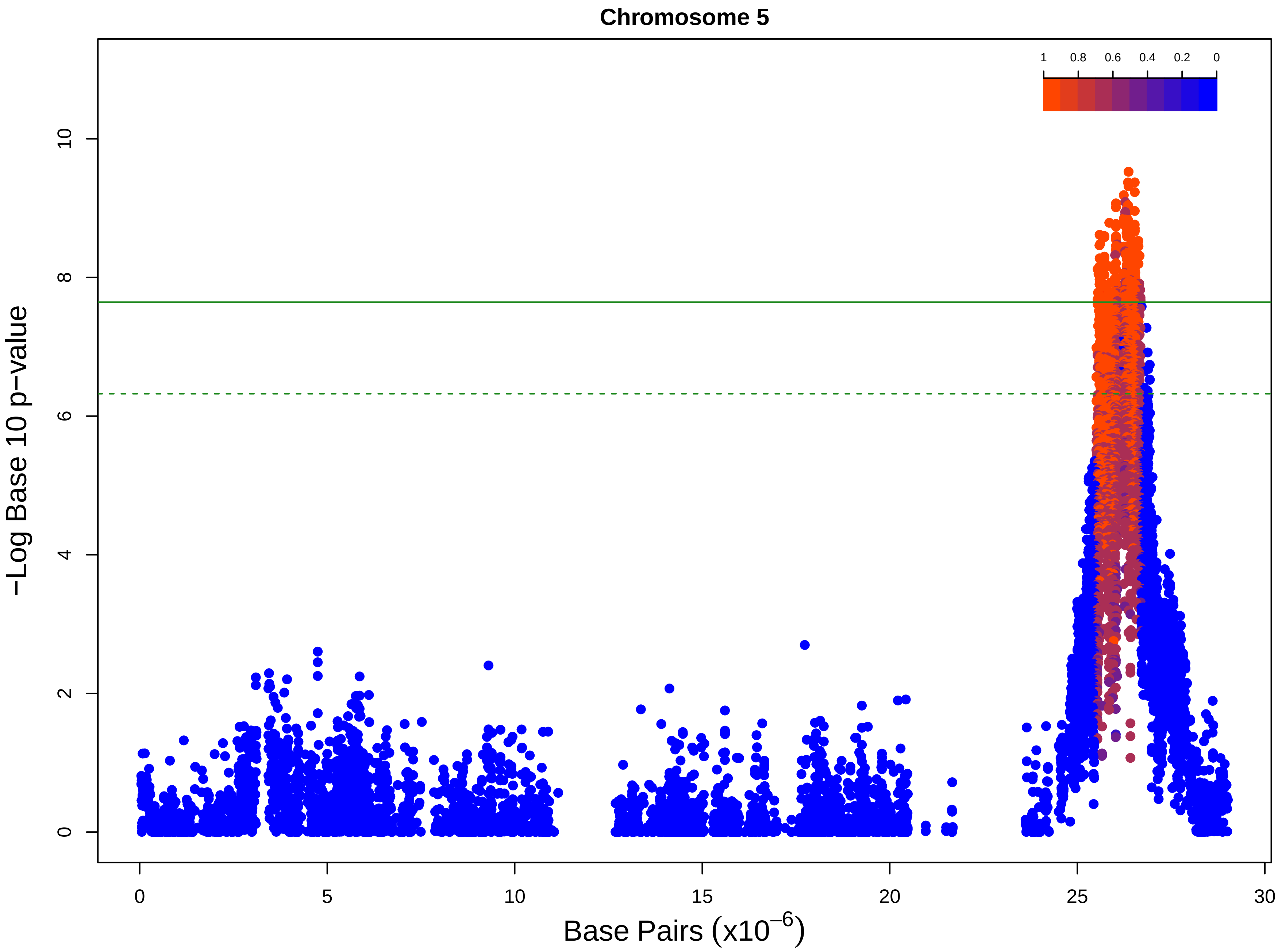

Chromosome 5
1
0.8
0.6
0.4
0.2
0
−Log Base 10 p−value
0
2
4
6
8
10
0
5
10
15
20
25
30
Base Pairs (x10$^{-6}$)

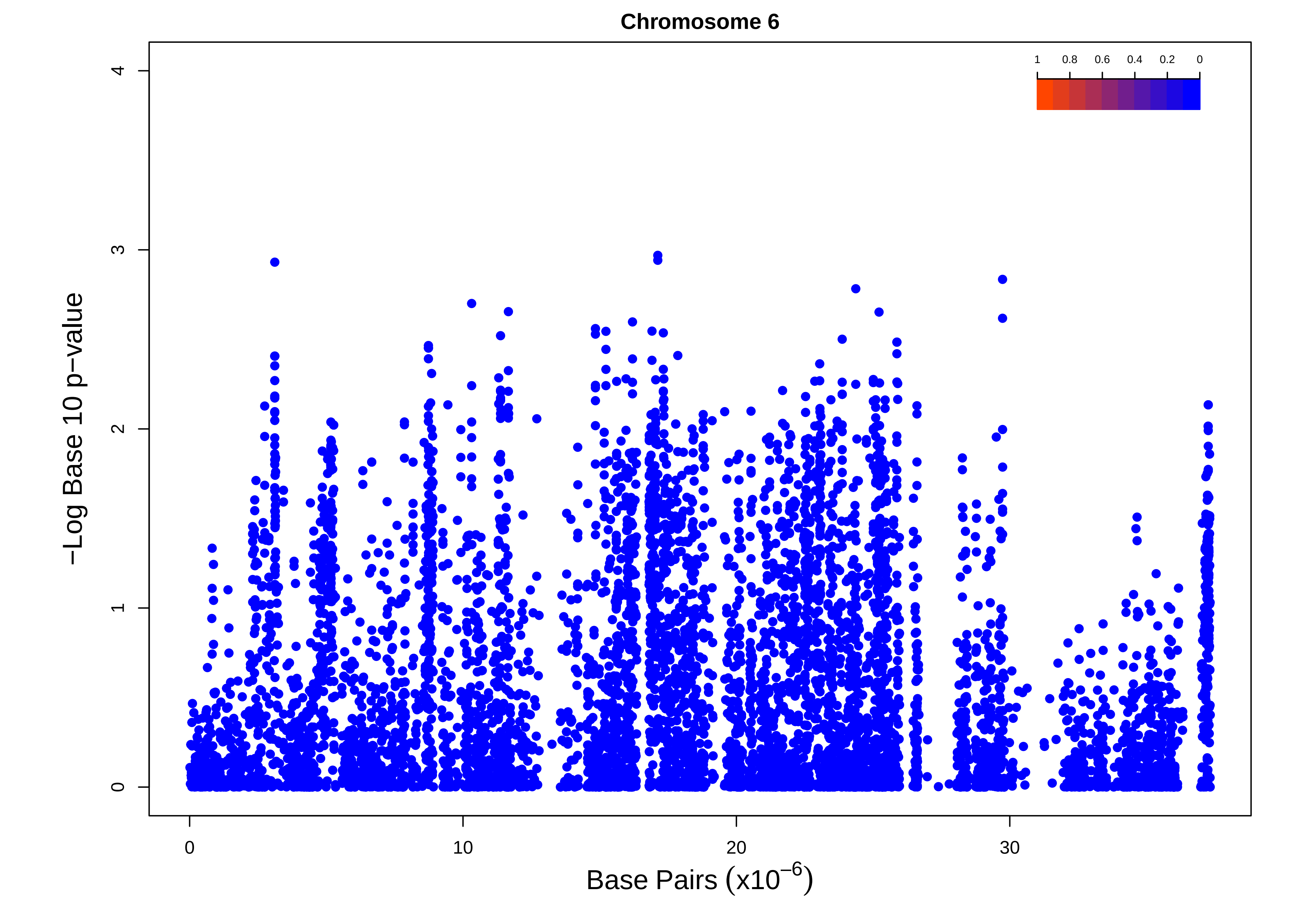
Chromosome 6
−Log Base 10 p−value
Base Pairs (x10^−6)
0
10
20
30
0
1
2
3
4
1
0.8
0.6
0.4
0.2
0

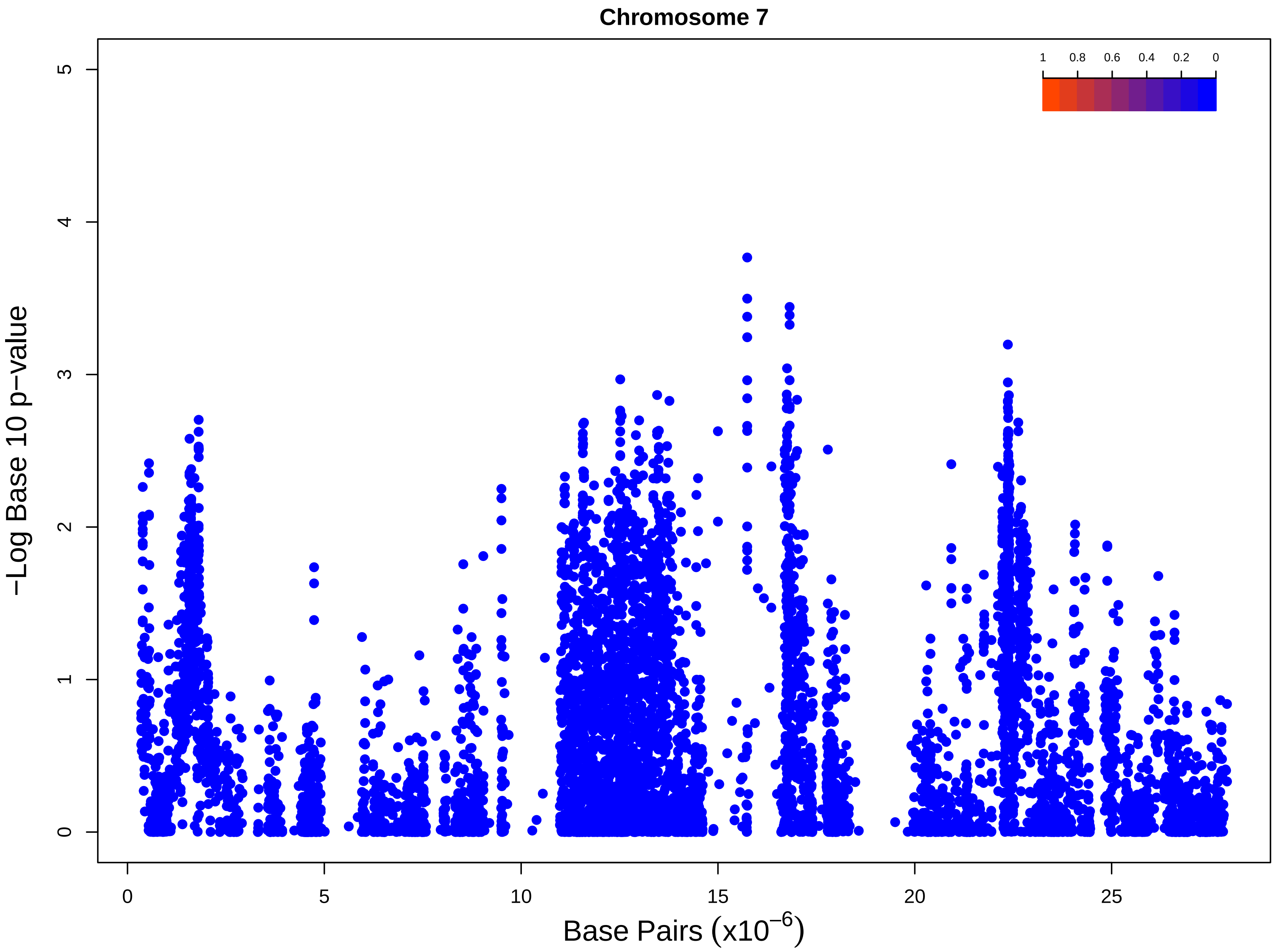
Chromosome 7
1
0.8
0.6
0.4
0.2
0
−Log Base 10 p−value
0
1
2
3
4
5
0
5
10
15
20
25
Base Pairs (x10^−6)

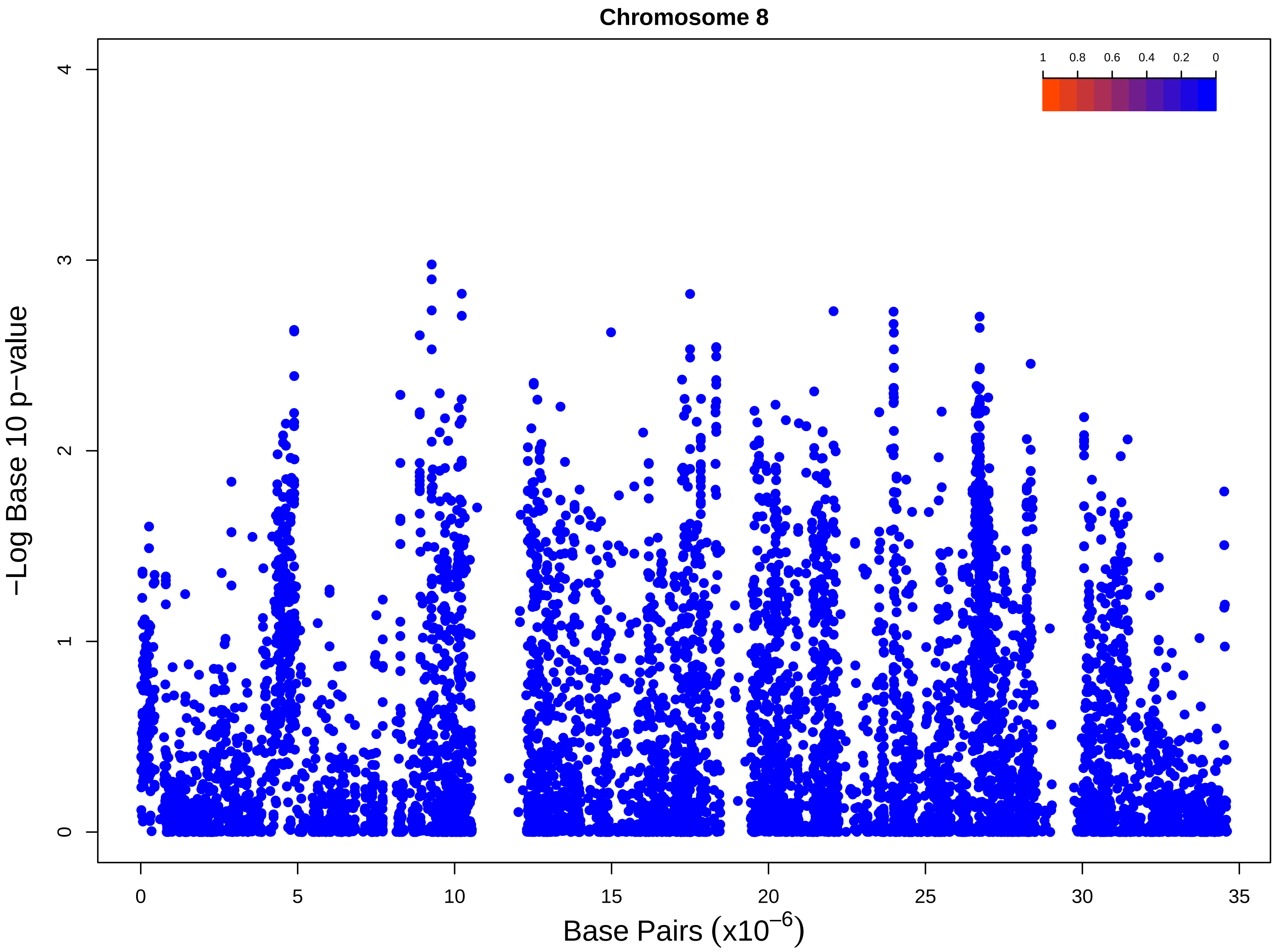

Chromosome 8
1
0.8
0.6
0.4
0.2
0
−Log Base 10 p−value
0
1
2
3
4
0
5
10
15
20
25
30
35
Base Pairs (x10^−6)

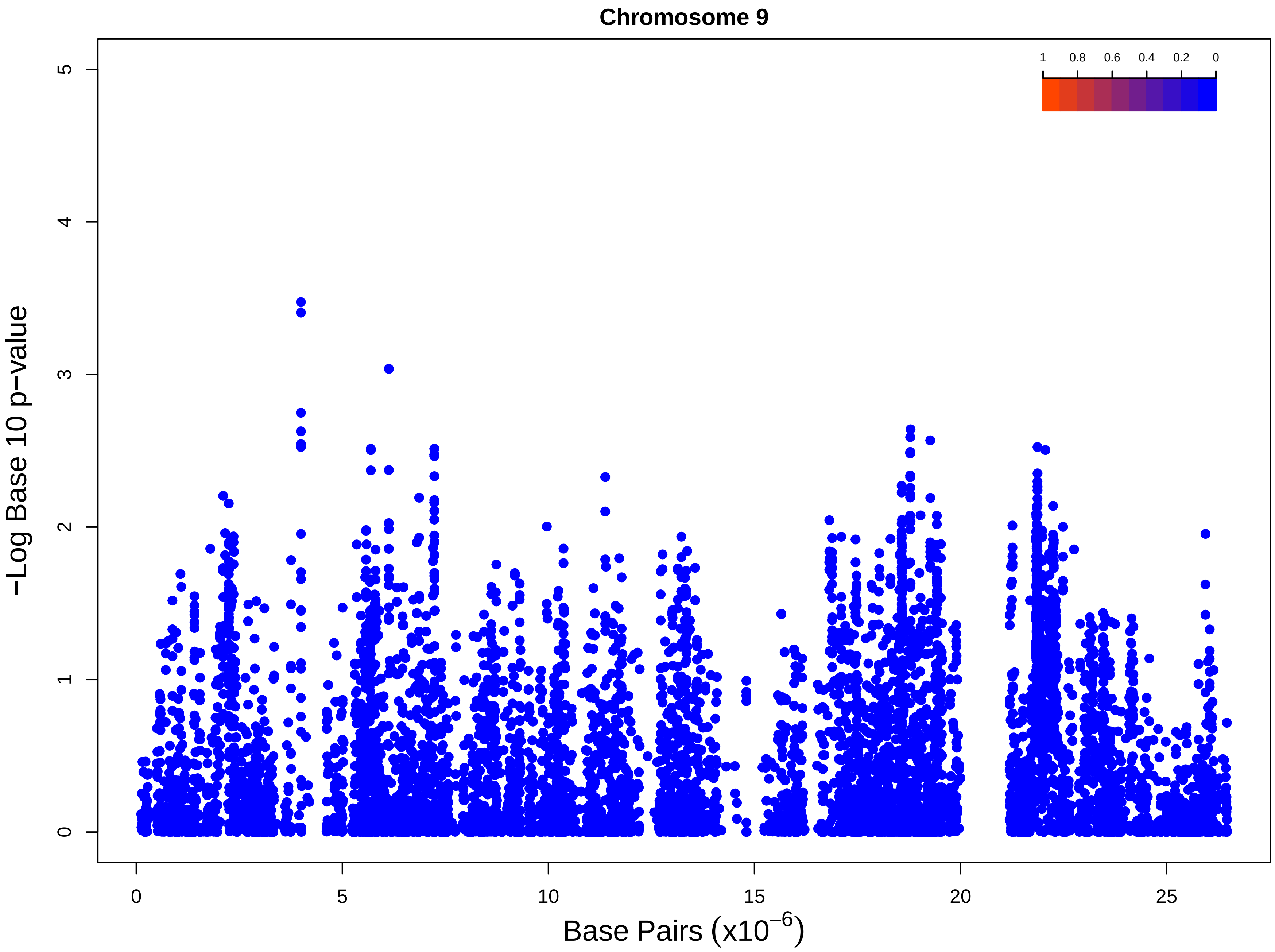
Chromosome 9
−Log Base 10 p−value
0
1
2
3
4
5
0
5
10
15
20
25
Base Pairs (x10^−6)
1
0.8
0.6
0.4
0.2
0

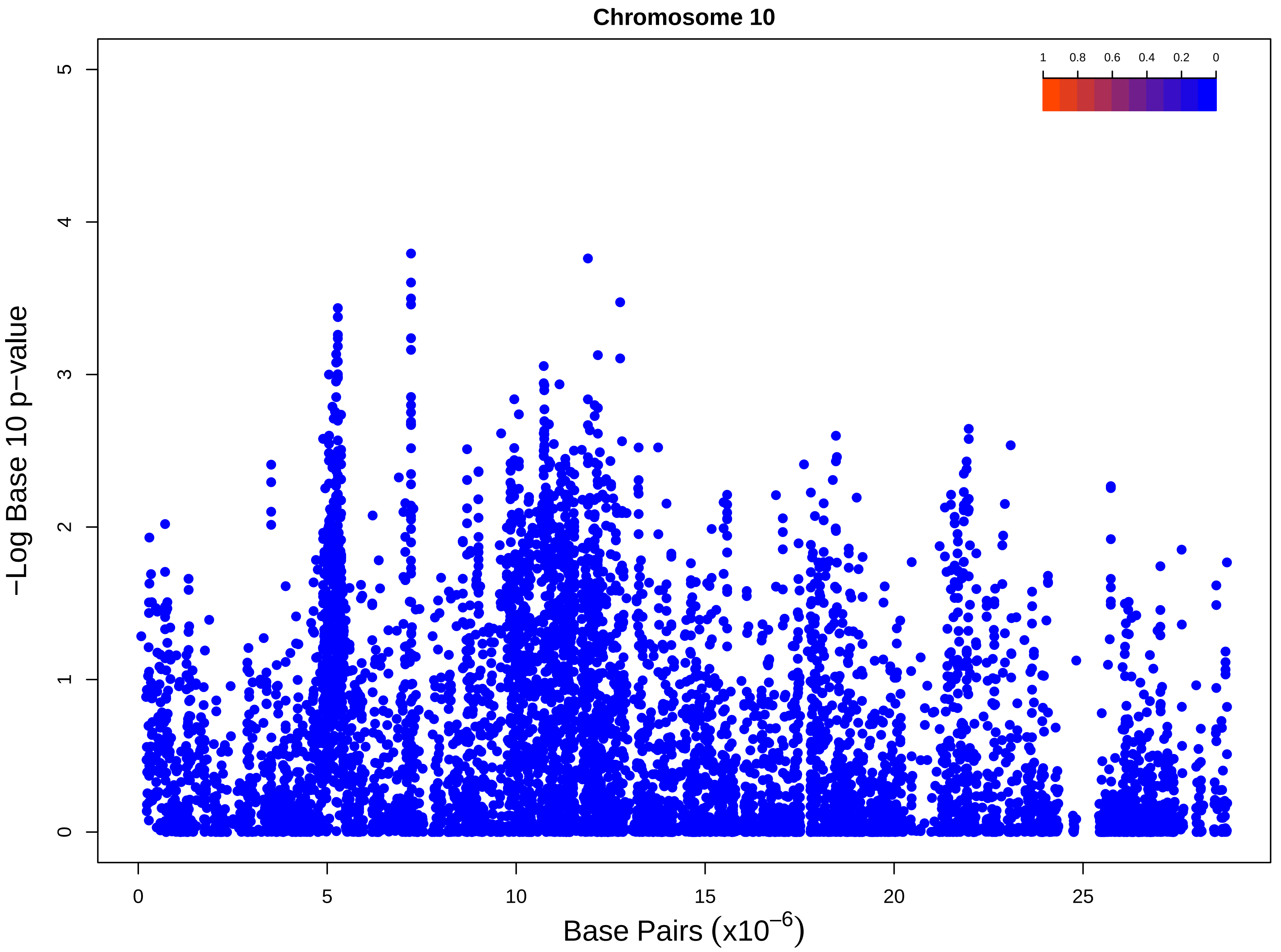

Chromosome 10
1
0.8
0.6
0.4
0.2
0
−Log Base 10 p−value
0
1
2
3
4
5
0
5
10
15
20
25
Base Pairs (x10^−6)

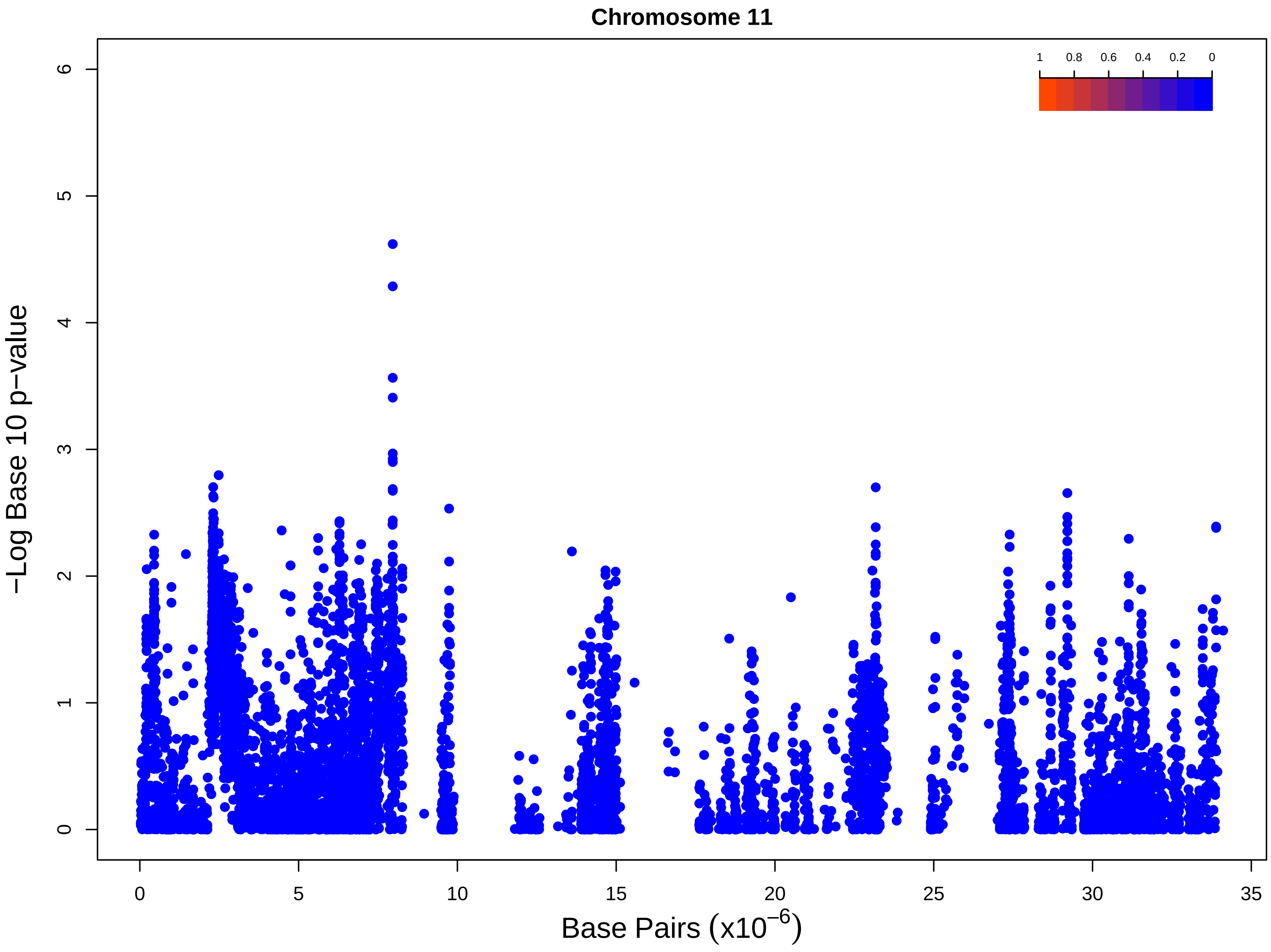
Chromosome 11
1
0.8
0.6
0.4
0.2
0
−Log Base 10 p−value
0
1
2
3
4
5
6
0
5
10
15
20
25
30
35
Base Pairs (x10^−6)

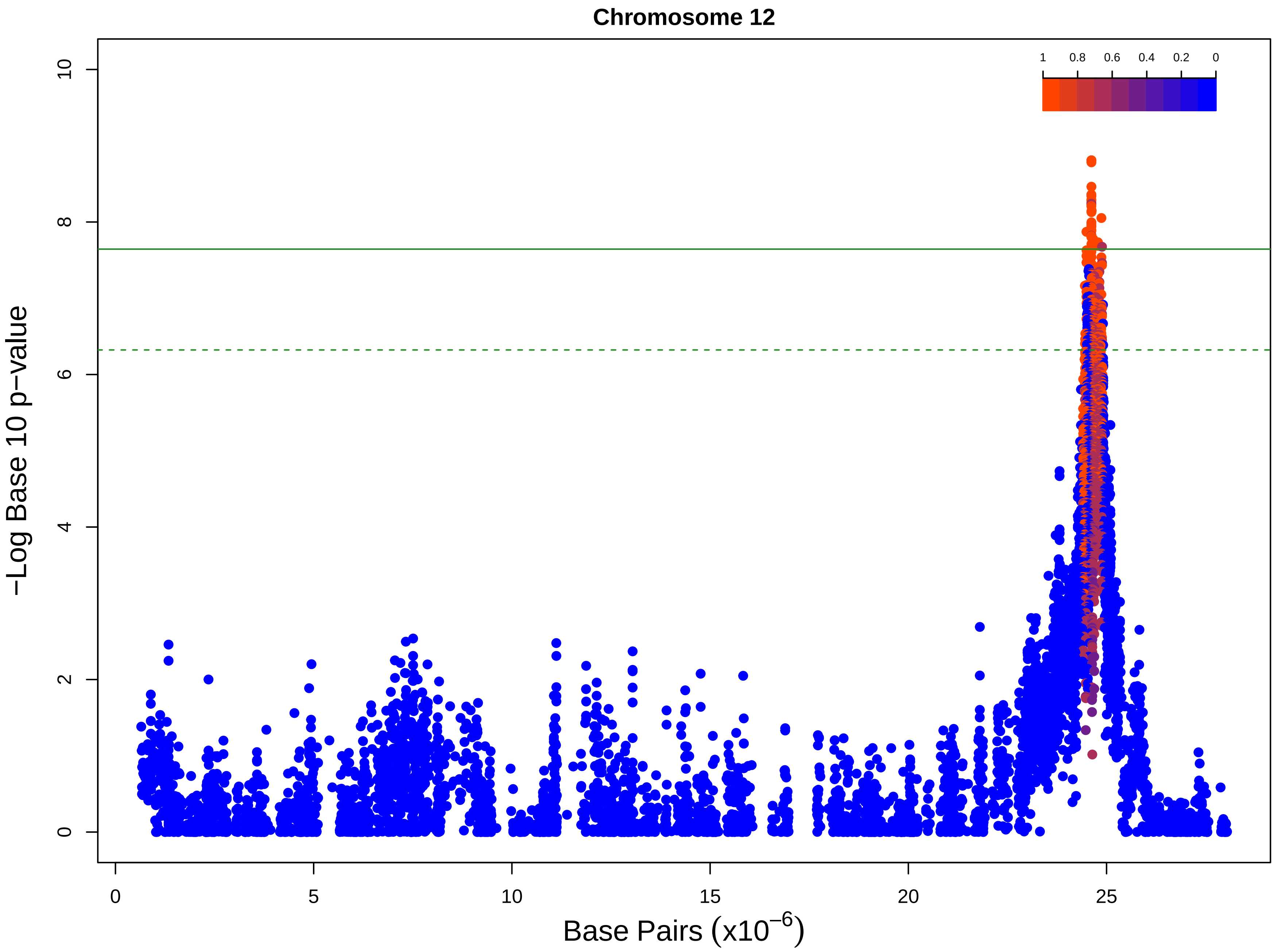

Chromosome 12
1
0.8
0.6
0.4
0.2
0
−Log Base 10 p−value
0
2
4
6
8
10
0
5
10
15
20
25
Base Pairs ($x10^{-6}$)
