## Supplementary figures and images for "Graph-based pangenomics maximizes genotyping density and reveals structural impacts on fungal resistance"

### combinedDotplot.png

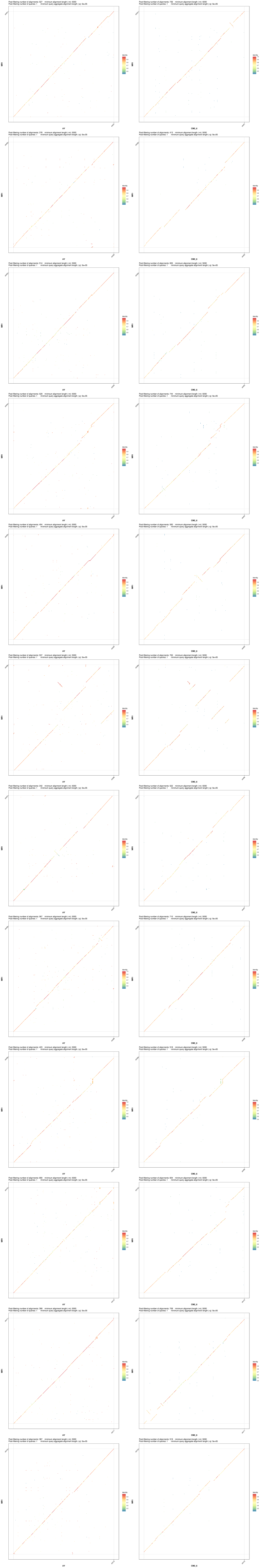

### fom2_protein_alignment.pdf

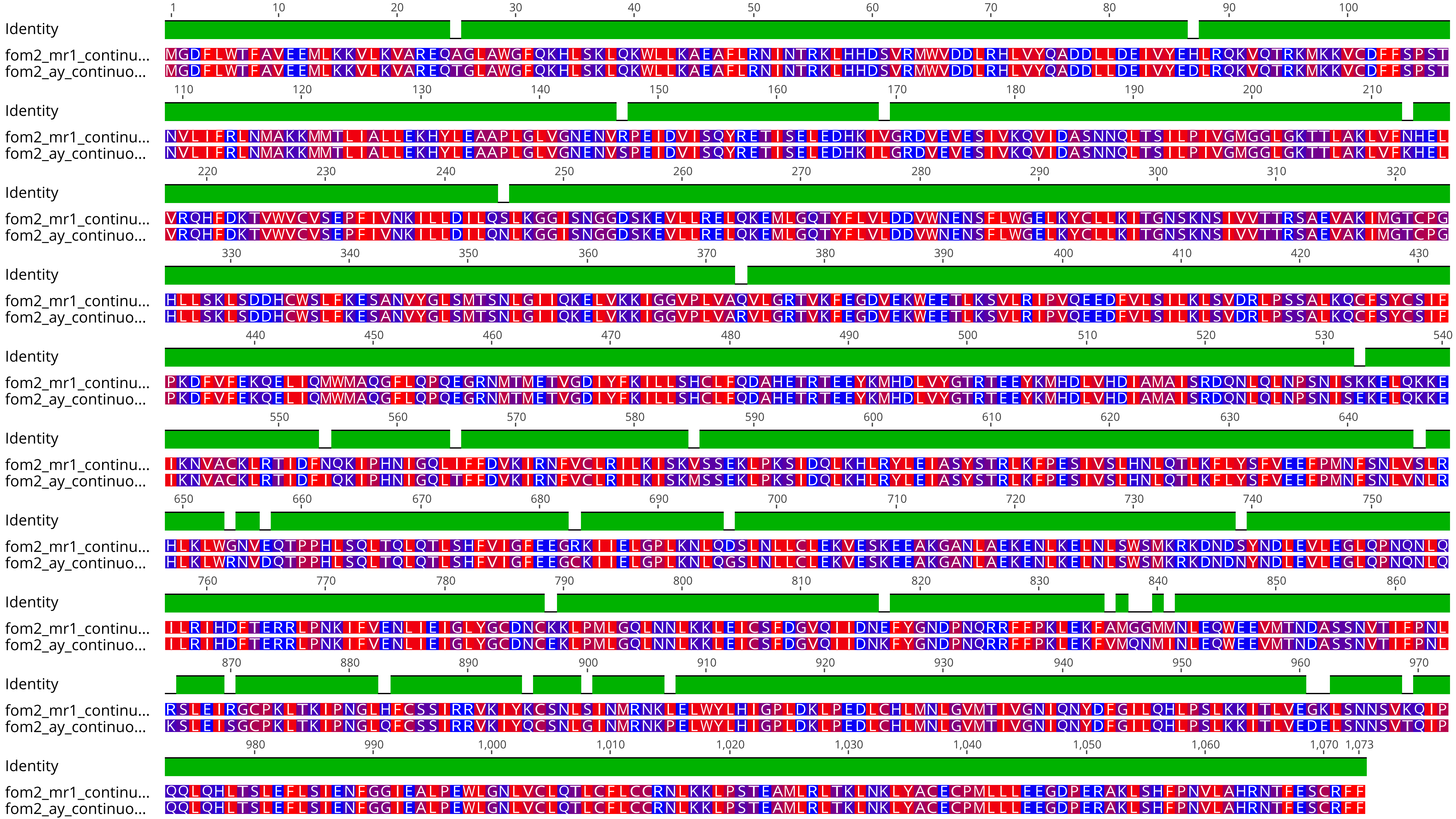

### fusAssay3Lines.jpg

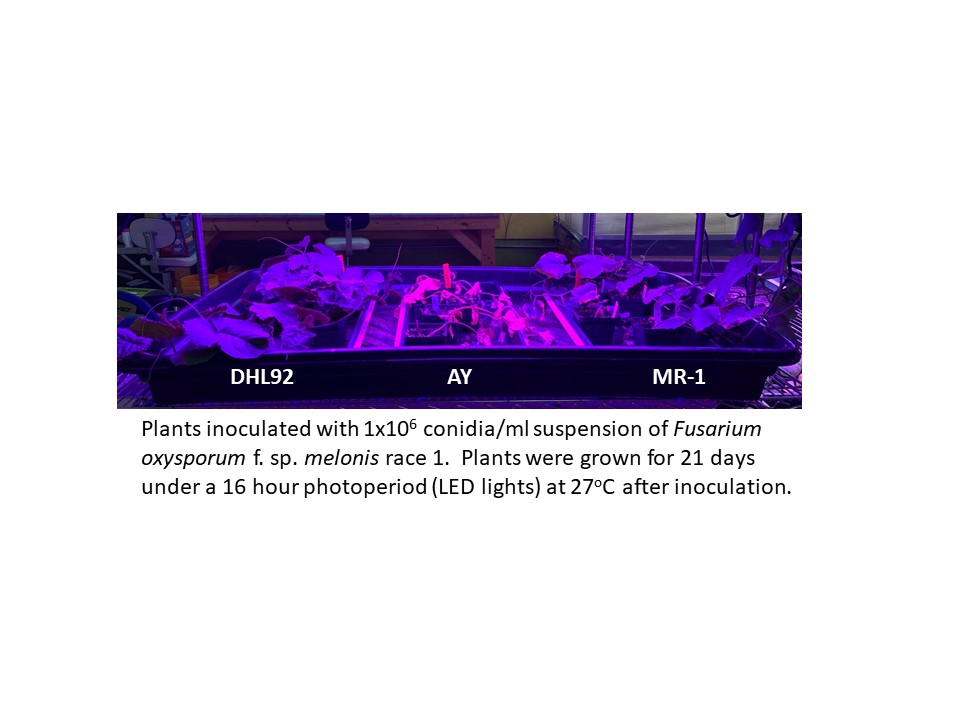

### uniqueNodes.venn.pdf

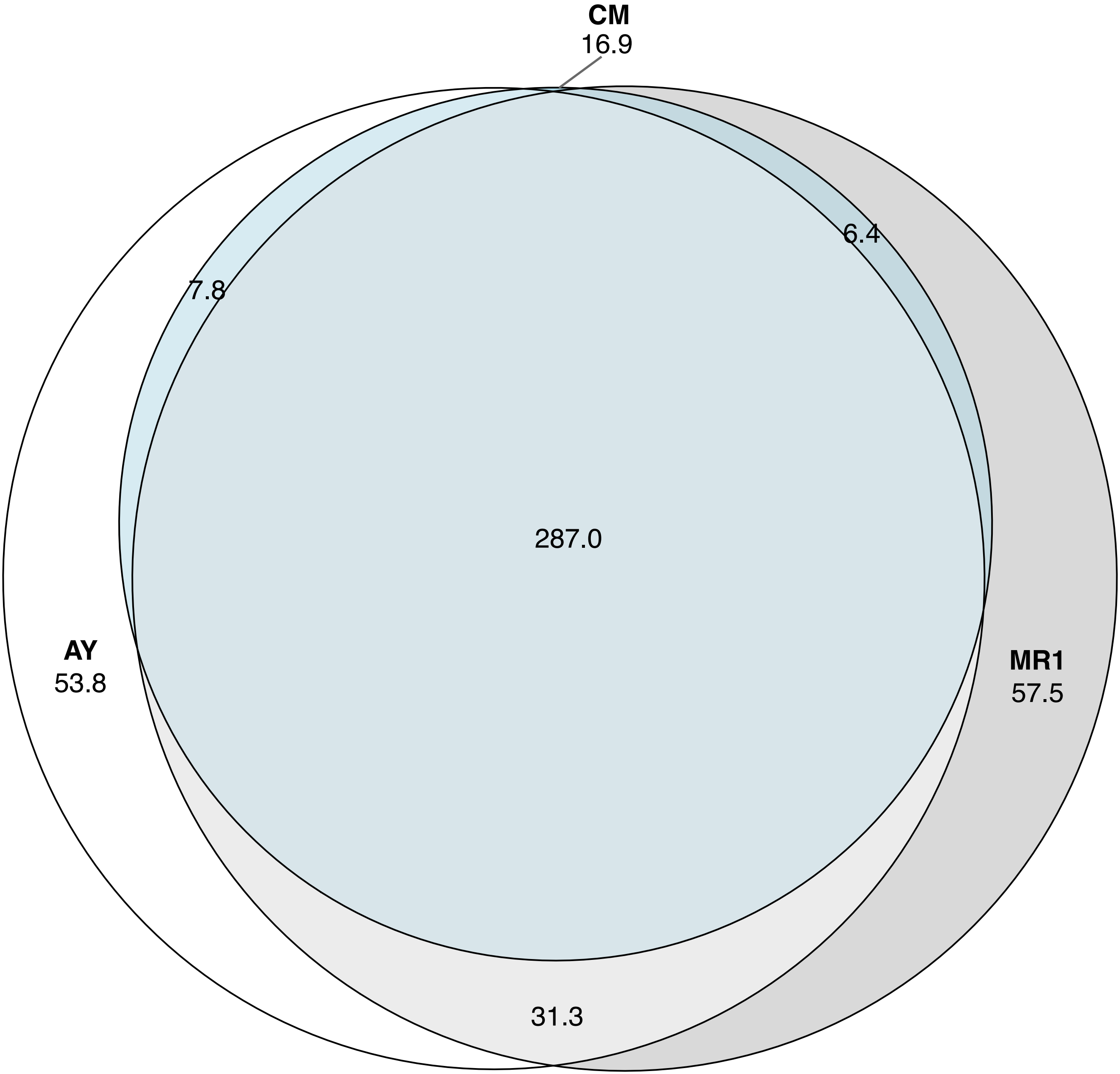

### variantDensity.pdf

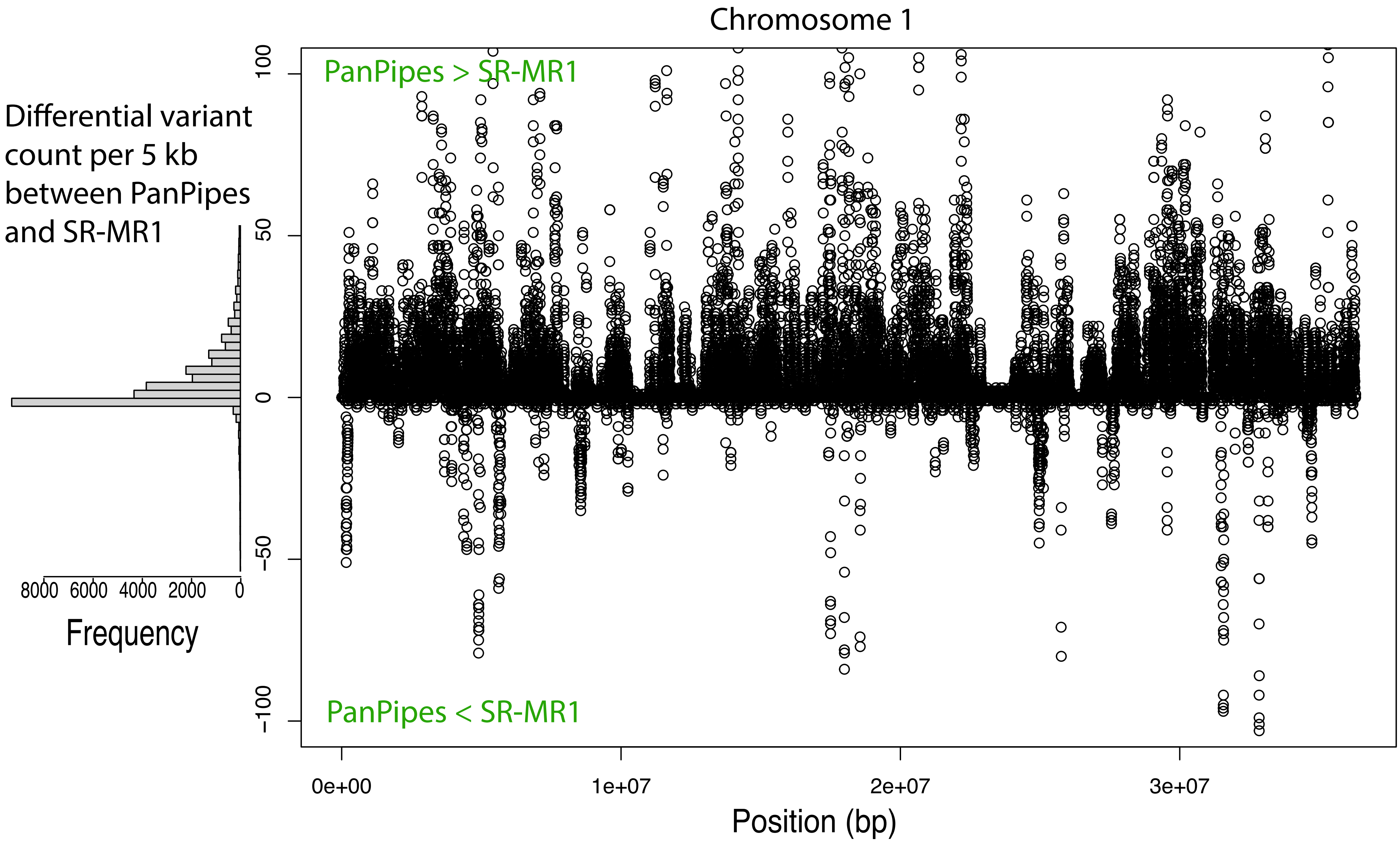
